# A *Plasmodium falciparum* lysophospholipase regulates fatty acid acquisition for membrane biogenesis to enable schizogonic asexual division

**DOI:** 10.1101/2021.07.01.450682

**Authors:** Pradeep Kumar Sheokand, Yoshiki Yamaryo-Botté, Vandana Thakur, Mudassir M. Banday, Mohd Asad, Cyrille Y. Botté, Asif Mohmmed

## Abstract

Phospholipid metabolism is crucial for membrane biogenesis and homeostasis during the intracellular life cycle of *Plasmodium falciparum.* To generate large amounts of phospholipids required during blood stages, the parasite massively scavenge, recycle and reassemble host lipids. *P. falciparum* possesses an unusual large number of lysophospholipases. However, their functional roles and importance remain to be elucidated. Here, we functionally characterized one of *P. falciparum* lysophospholipase (*Pf*LPL3) (Gene ID PF3D7_1476800), to reveal its critical role in parasite propagation during asexual blood stages. We generated a transgenic parasite line using GFP-*glm*S C-terminal tagging approach, for localization as well as inducible knockdown of *Pf*LPL3. *Pf*LPL3 displayed a dynamic localization throughout asexual stages, mainly localizing in the host parasite interface: parasitophorous vacuole space and expanding into the tubulovesicular network within the host cell. Inducible knock-down of *Pf*LPL3 hindered normal intraerythrocytic cycle, specifically causing disruption in parasite development from trophozoites to schizont, as well as reduction in number of merozoites progenies. Thus, down-regulation of *Pf*LPL3 significantly inhibited parasite growth suggesting its critical role for proper parasite propagation during blood stages. Detailed lipidomic analyses showed that *Pf*LPL3 generates fatty-acids for the synthesis of neutral lipids DAG and TAG, whilst controlling the timely synthesis of phospholipids that are crucial for membrane biogenesis required for merozoite development during asexual cycle. Setting up an *in vitro* activity based screening of ‘Malaria Box’ allowed identification of specific inhibitors of *Pf*LPL3 having potent parasitical efficacies. These compounds are pertinent both as anti-malarial drug candidates and chemical tools specifically targeting membrane biogenesis during asexual blood stages.

## Introduction

Malaria is a vector-transmitted parasitic disease, which remains a major medical problem in tropical and sub-tropical areas of the world. Malaria leads to ~500,000 to 1 million deaths globally every year (1) WHO, (2). In absence of efficient vaccine and due to rapid spread of drug resistant parasites strains, there is urgent need to identify of new drug-targets and development of new drugs against the disease. Understanding of key metabolic pathways in the parasite, which are essential for parasite survival, is a crucial pre-requisite to identify unique and potent targets in the parasites. The pathological symptoms of malaria are caused by repeated blood stage asexual cycles of the parasite in host erythrocytes. Different parasite development stages during the asexual life cycle involve extensive coordinated lipid synthesis, modification of host cell membrane, and biogenesis membrane needed for parasite division and propagation (3). This high demand of lipids/phospholipids required for extensive membrane synthesis is fulfilled by: scavenging from the host milieu, recycling and modifying host lipids, *de novo* synthesis and re-shuffling of those through parasite lipid synthesis machinery (4, 5). The high demand in lipids by the parasite is perfectly illustrated in *P. falciparum* asexual blood stages where lipid content increases by 200-300% through the acquisition and synthesis of a large number of lipid species for growth and proliferation (3, 6). Phospholipids are the major structural components of parasite membranes and are used to make new daughter cells. PC is the most abundant phospholipid in *Plasmodium* membranes during both liver stages and blood stages, attesting its importance for parasite division and propagation (4, 7). The parasite possesses the typical eukaryotic *de novo* phospholipid synthesis machineries to generate most (lyso)phospholipids classes and their precursors from fatty acids (FA), and polar heads, i.e. the so-called Kennedy pathway and the CDP-DAG pathway. *P. falciparum* blood stages notably relies on the scavenging of lysophosphatidylcholine (LPC), especially its polar head phosphocholine, to fuel this *de novo* synthesis of PC and maintain high asexual division rates (8, 9). Absence of LPC in the extracellular media drives parasite sexual differentiation into gametocytes, likely by lack of PC used as building material for active asexual division. Since LPC plays an important role in parasite development and differentiation, enzymes involved in its hydrolysis are considered as attractive targets for antimalarial development. The catabolism of lipid metabolites from the host must involve phospholipases to manipulate the scavenged lipids and generate the proper precursors (i.e. FA and polar heads). Indeed, parasite contains a high number of phospholipases, harbouring α/β hydrolase domain, including a family of lysophospholipase (LPL). The LPLs play key role recycling of lipids by hydrolysis of acyl chains from lysophospholipids, which are the intermediated in metabolism of membrane phospholipids.

In the present study, we have functionally characterized one of the *P. falciparum* lysophospholipase, *Pf*LPL3 (Gene ID PF3D7_1476800) that is essential for blood stage schizogonic division through recycling of host lipids. We used a GFP-*glm*S ribozyme approach (10), which allowed the endogenous localization of the protein as well as inducible-transient down-regulation during the parasite blood stages. Confocal analyses showed that *Pf*LPL3 has a dynamic localization at interfaces between the parasite and the host cell, at the parasitophorous vacuole, as well as extending into the tubulovesicular network space within the host erythrocytes, more specifically during late-trophozoite and schizont stages. Transient knock-down of *Pf*LPL3 caused dramatic hindrance in schizont development as well as significant decrease in number of merozoites progenies generated in each schizont. To determine the function of *Pf*LPL3 for merozoite formation and parasite survival, we performed state-of-the-art lipidomic analyses, which showed that disruption of *Pf*LPL3 induced drastic changes in parasite lipid composition and homeostasis, impacting fatty acids (FA), phospholipids and neutral lipid contents. Specifically, lipidomic analyses revealed that *Pf*LPL3 generates free FA (FFA) that are normally fuelling the DAG-TAG synthesis pathway for lipid storage, and in turn regulate the active synthesis of phospholipids, which are essentially required for membrane biogenesis of merozoites during schizogony. We developed an *in vitro* activity assay using recombinant *Pf*LPL3 that confirmed LPL activity for the enzyme. Using this activity assay, we screened the existing MMV “Malaria Box” compounds chemo library. We were able to identify two specific inhibitors of *Pf*LPL3 enzymatic activities with potent parasitical efficacies (<1.0 μM range). Importantly, treatment of blood stage parasites phenocopied the disruption of *Pf*LPL3 in the parasite, confirming their selectivity for *Pf*LPL3. Based on the importance of *Pf*LPL3 for blood stage parasite development, these compounds could be further developed as new antimalarials and/or chemical tools specifically targeting lipid homeostasis machinery in malaria parasite.

## Results

### Endogenous tagging of *pfLPL3* gene and localization of fusion protein in transgenic parasites

We tagged the endogenous *Pf*LPL3 for physiological localization studies as well as to study its functional essentially by conditional knock-down strategy. We used GFP-*glm*S ribozyme system (10) for C-terminal tagging of the native *pfLPL3* gene, so that the fusion protein gets expressed under the control of native promoter (Fig. 1A). Integration of the tag at the C-terminus of the gene was confirmed in selected clonal parasite population using PCR-based analyses (Fig. 1B). Expression of fusion protein of calculated size of ~70 kDa was detected by western blot analysis using anti-GFP antibody (Fig. 1C). This band was not detected in wild type 3D7 parasites. These transgenic parasites, *Pf*LPL3_GFP_*glm*S, were studied to determine the physiological localization of the *Pf*LPL3-GFP fusion protein by fluorescence and confocal microscopy approaches. The chimeric protein was expressed throughout all stages of the asexual blood life cycle of the parasite (Fig. 1D). In early development stages, ring and mid-trophozoite stages, GFP fluorescence was mainly observed in the cytosol. As parasites matured into late-trophozoite and schizonts stages, the GFP labelling was observed at the parasite periphery in the parasitophorous vacuole (PV) region. Intriguingly, during the late trophozoite stages, *Pf*LPL3 was also present in the parasite extension of host-parasite interface structure: within the tubulovesicular networks (TVN) (Fig. 1D; Supplementary Fig. S2A). A 3D reconstruction of trophozoite stage parasite images clearly confirmed presence of GFP staining around the parasite periphery and in the TVN (Fig. 2A). To find out if the *Pf*LPL3 is present outside the parasite plasma membrane, infected RBCs from transgenic parasite culture were fractionated by saponin lysis. *Pf*LPL3 was detected in the supernatant as well as in the pellet fraction by western blot analysis, which confirmed the presence of *Pf*LPL3 in the PV as well as in parasite cytosol (Fig. 2B), correlating with the fluorescence microscopy results.

**Figure 1:**
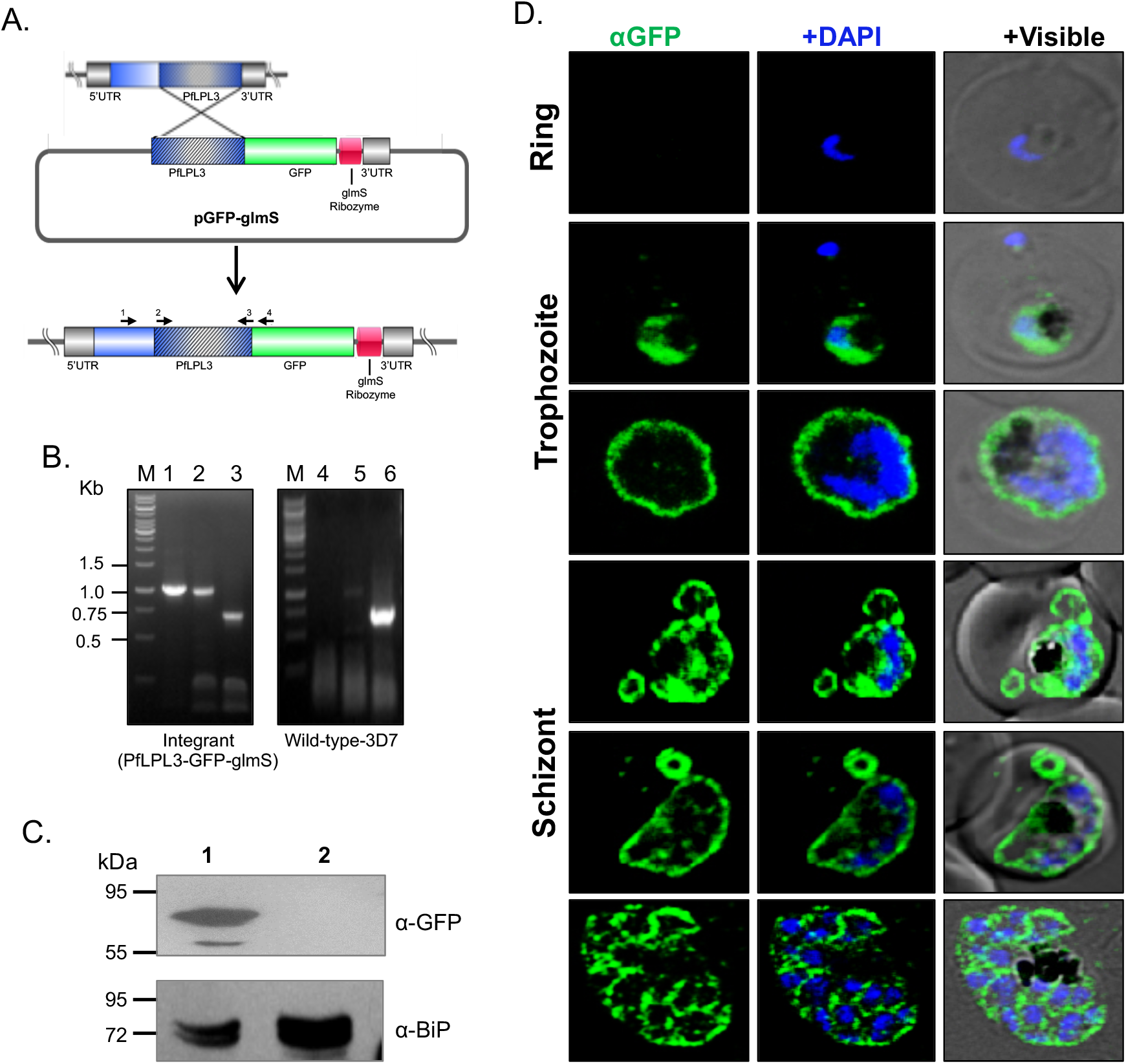
Generation of *pf*LPL3-GFP-*glm*S transgenic parasite line: (A) A schematic representation of single crossover homologous recombination showing the integration of the LPL3-GFP-*glm*S plasmid at the C-terminus of the endogenous *pflpl3* gene. (B) PCR based analyses of the selected clonal parasite culture and wild type 3D7 parasite lines, locations of primers used are marked in the schematic. Lane 1 and 4 (primers 1078A and 1234A) showing amplification only in the integrants; lane 2 and 4 (primers 1222A and 1234A) show amplification in integrants or episome parasites; lane 3 and 6 (primers number 1222A and 1223A) show amplification in both integrants and the wild-type line. (C) Western blot of transgenic parasites probed with anti-GFP antibody. The fusion protein band was detected in the transgenic parasites, *Pf*LPL3-GFP-*glm*S only (lane 1) and not in the 3D7 parent parasite (lane 2). Blot ran in paralled and probed with anti-BiP was used as loading control. (D) Fluorescent microscopic images of live transgenic parasites at ring, trophozoite, and schizont stages. The parasite nuclei were stained with DAPI and parasites were visualized by confocal laser scanning microscope. In young parasites, the fusion protein was observed in the cytosol of the parasite, but as the parasite matures, the fluorescence was observed towards the periphery of the parasite.

**Figure 2:**
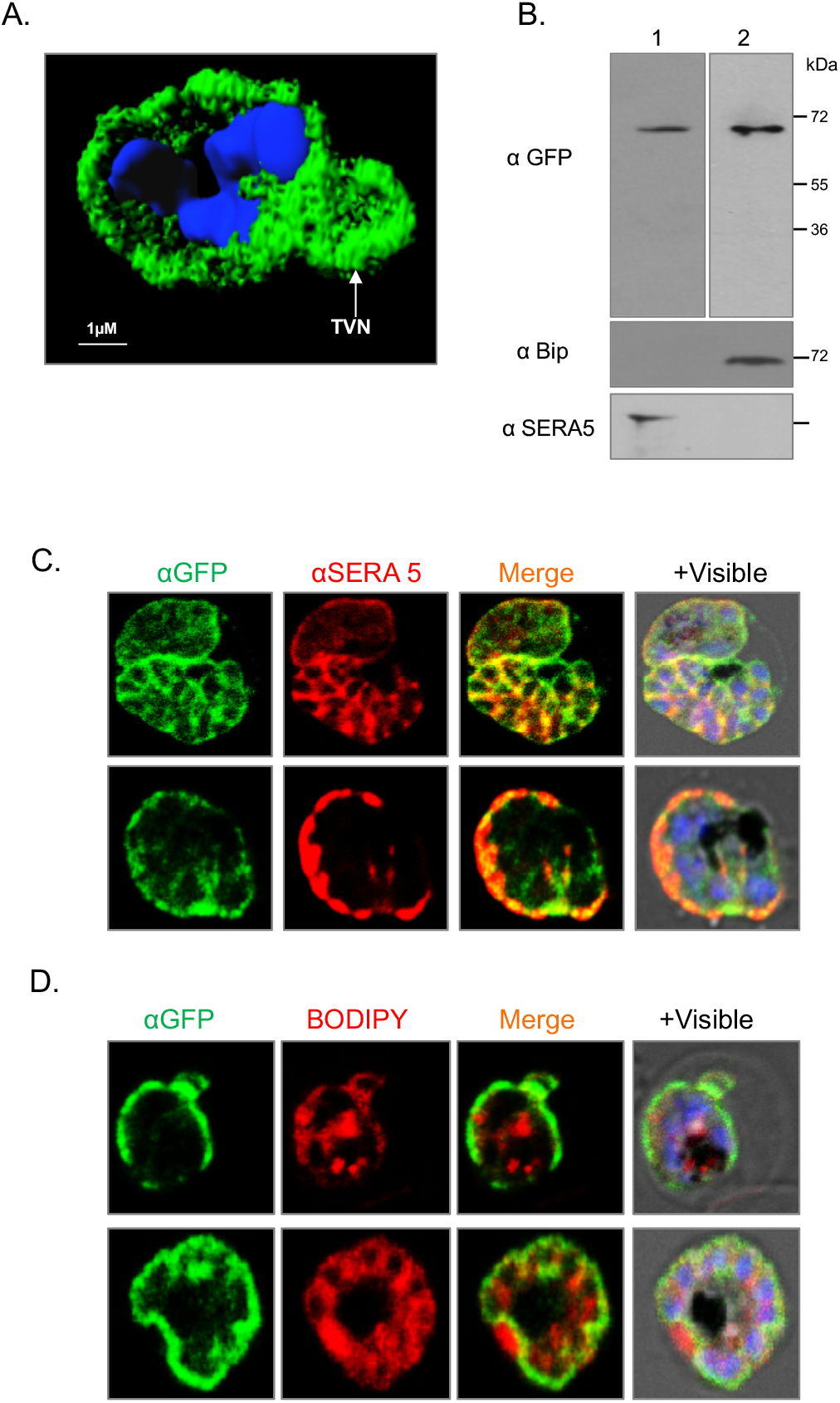
Sub-cellular localization of *Pf*LPL3 in parasitophorous vacuole (PV) and tubulovesicular network: (A) The 3D image constructed by using series of Z-stack images of the transgenic parasites with IMARIS 7.0, shows the GFP-fusion protein present in the PV and tubulovesicular network (TVN) region. (B) Western blot of infected erythrocytes fractions after saponin lysis showing that the *Pf*LPL3-GFP fusion protein is present in soluble fraction representing content of parsitophorous vacuole (PV) and host cytosol; blot ran in parallel and probed with anti-BiP and anti-SERA5 antibody were used as a negative and positive controls respectively. (C) Fluorescent microscopic images of trangenic parasites immustained with anti-SERA5 antibody showing colocalisation of *Pf*LPL3 with SERA5 in PV. (D) Fluorescent microscopic images of transgenic parasites stained with mambrane probe BODIPY-TR ceramide, showing GFP fusion protein at the parasite periphery in late parsite stages. Parasite nuclei were stained with DAPI and images were acquired by confocal laser scanning microscope.

To further investigate these localization patterns, membrane labelling and immunostaining studies were carried out. The GFP tagged parasites were co-stained with the lipid membrane probe BODIPY-TR ceramide, which stains all membranous structures. The BODIPY-Ceramide probe labeled parasite periphery, which include plasma membrane and PV membrane (PVM) (Fig. 2C). In addition, the tubulovesicular extension of the PVM were observed in the erythrocyte cytosol for a section of the late trophozoite/schizont stage parasite population (Fig. 2B, C, D, 1D). The *Pf*LPL3-GFP labelling showed overlap with BODIPY, in the PV region as well as in the TVN (Figure 2C and S2B). Further, co-labeling of transgenic parasites with one of the PV-resident protein, SERA5 (11), confirmed the localization of *Pf*LPL3-GFP protein in the PV (Fig. 2D and S2C).

### Selective degradation of *Pf*LPL3 inhibits parasite growth

To understand the functional significance of *Pf*LPL3 and its possible involvement in lipid metabolism, we utilized the *Pf*LPL3_GFP_*glm*S parasite line for inducible knock-down of *Pf*LPL3 expression using *glm*S ribozyme. In presence of glucosamine (GlcN), the *glm*S-ribozyme cleaves itself, which in turn leads to degradation of the associated *Pf*LPL3 mRNA. Selective knock-down of *Pf*LPL3 protein was assessed in transgenic parasite treated with different GlcN concentrations (1.25mM, 2.5mM, 5mM and 10mM). The parasites showed GlcN concentration dependent reduction in *Pf*LPL3 levels (60-90%) in treated sets (Fig. S3A). To avoid any potential GlcN-mediated toxicity observed at high GlcN concentrations, all further experiments were carried out using 1.25mM of GlcN. To study the effect of the inducible knock-down of *Pf*LPL3 (*Pf*LPL3-iKD) on parasite growth, total parasitemia was determined at different time points (24h, 36h, 48h) after GlcN treatment. The *Pf*LPL3-iKD set showed ~70% inhibition of parasite growth compared to untreated parasites (Fig. 3A). There was no deleterious effect of 1.25mM GlcN on the growth of the wild type parasite line.

**Figure 3:**
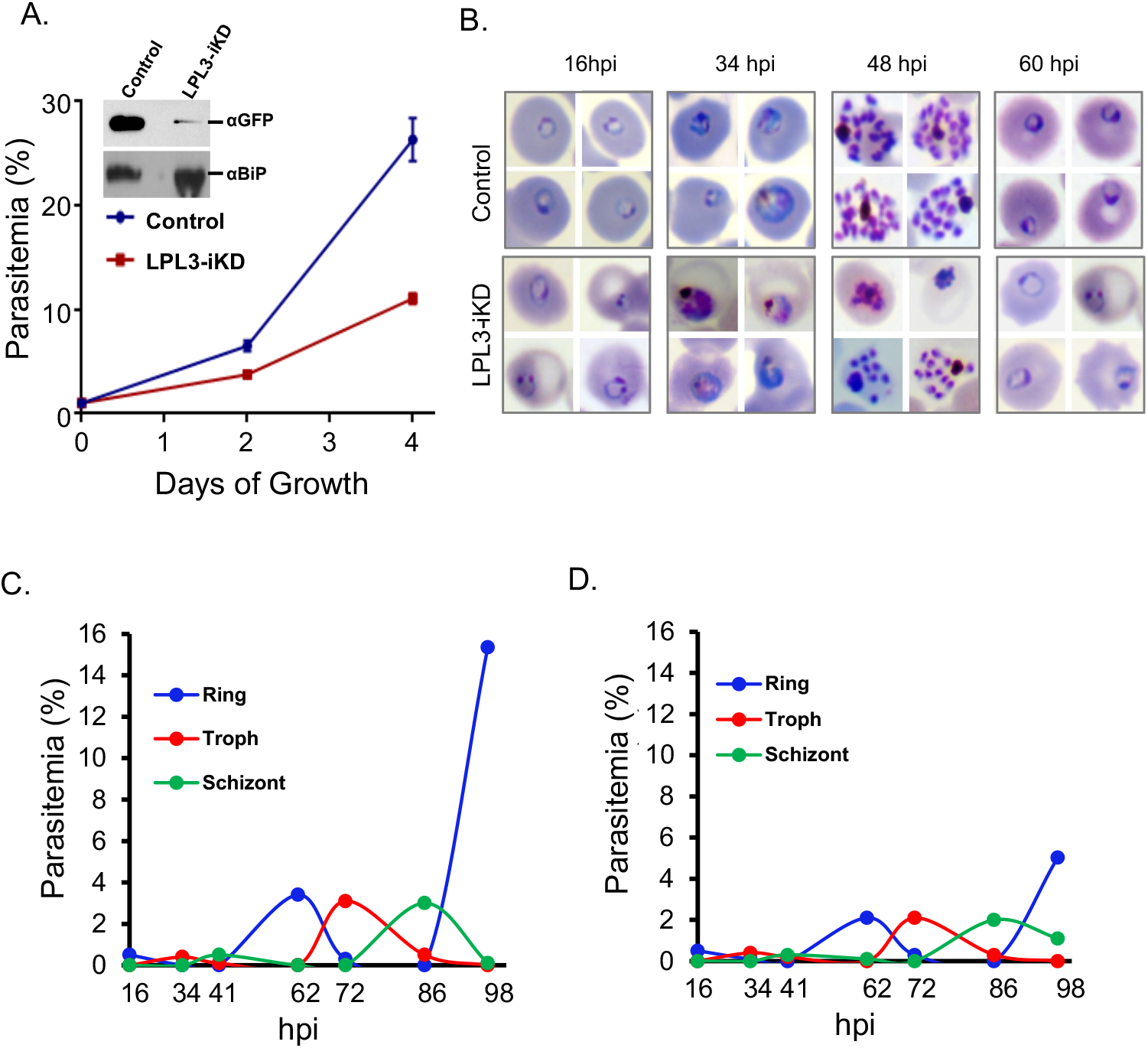
Inducible knock-down of *Pf*LPL3 hampers parasite growth and survival: (A) Graph showing percetage parasitemia in *Pf*LPL3-GFP-*glm*S transgenic parasite cultures grown with 2.5 mM glucosamine (iKD and control), as estimated by new ring stage parasites after 2 and 4 days of growth. Western blot analysis showing reduction in the fusion protein in presence of glucosaime is shown in inset. (B) Giemsa stained images of parasites showing effect on parasite morphology at different time points (0-48h) for control and *Pf*LPL3-iKD sets. Parasite stage progression curve for (C) control and (D) *Pf*LPL3-iKD sets.

### *Pf*LPL3 is important for parasite intracellular development and crucial for normal schizogonic division

To study the effect of selective degradation of *Pf*LPL3 on parasite development and morphology, the *Pf*LPL3-iKD set and control set were monitored after different time intervals (Fig. 3B, C and D). In control set, during each intra-erythrocytic cycle, the parasites usually develop from ring to trophozoites to mature schizonts, and subsequent merozoites released from these schizonts can invade new erythrocytes. Together, the intra-erythrocytic cycle can effectively increase the total parasitemia for about 5-6 fold/48h cycle in our culture conditions (Fig. 3A). In the *Pf*LPL3-iKD set, parasite growth could only increase up to two folds (Fig. 3). Morphological observations on parasite intracellular development showed no effect on development of ring stages in to trophozoite stages in *Pf*LPL3-iKD as compared to control set (Fig. 3B). However, aberrant development of late trophozoites and schizonts were observed in *Pf*LPL3-iKD, having smaller, aberrant, or sick-looking parasites compared to control (Fig. 3B); a large number of trophozoites were not able develop into schizonts, which caused ~50% reduction in number of schizonts as compared to control (Fig. 3C, D). Indeed, at 48 hpi (hours post invasion), a number of stressed parasites were observed in *Pf*LPL3-iKD set as compared to control set (Fig. 3B, C), which were not viable to develop into schizonts (Fig. 3D). Further, the *Pf*LPL3-iKD parasites were more specifically affected at the replicative fitness of these schizonts. During schizogony *P. falciparum* typically undergoes multiple rounds of nuclear divisions resulting in 16-32 merozoites. In the control set, majority of the schizonts were found to contain 16 or more merozoites (Fig. 4A). However, in the *Pf*LPL3-iKD parasite set, mean number of merozoites per schizonts was significantly reduced (Fig. 4A). This decrease in replicative fitness of schizonts resulted in ~25% decrease in total number of merozoites with respect to segmented schizonts (Fig. 4B). It is to be pointed out this effect is in addition to hampering of parasite development in the *Pf*LPL3-iKD set, and a number of parasites were developmentally stuck even before the segmental stages; therefore, only few of the parasite could develop into segmented schizonts, which also harbored reduced number of merozoites as compared to control. However, immunostaining of these schizonts with anti-MSP1 antibody showed that there was no defect in the merozoites membrane in the *Pf*LPL3-iKD set, which suggested that the merozoite generated had fully developed plasma membrane (Fig 4C). Overall, we show that *Pf*LPL3-iKD not only inhibited schizont development (>50%), but it also reduced replicative fitness of parasites (~25%), which resulted in >70% parasite growth inhibition.

**Figure 4:**
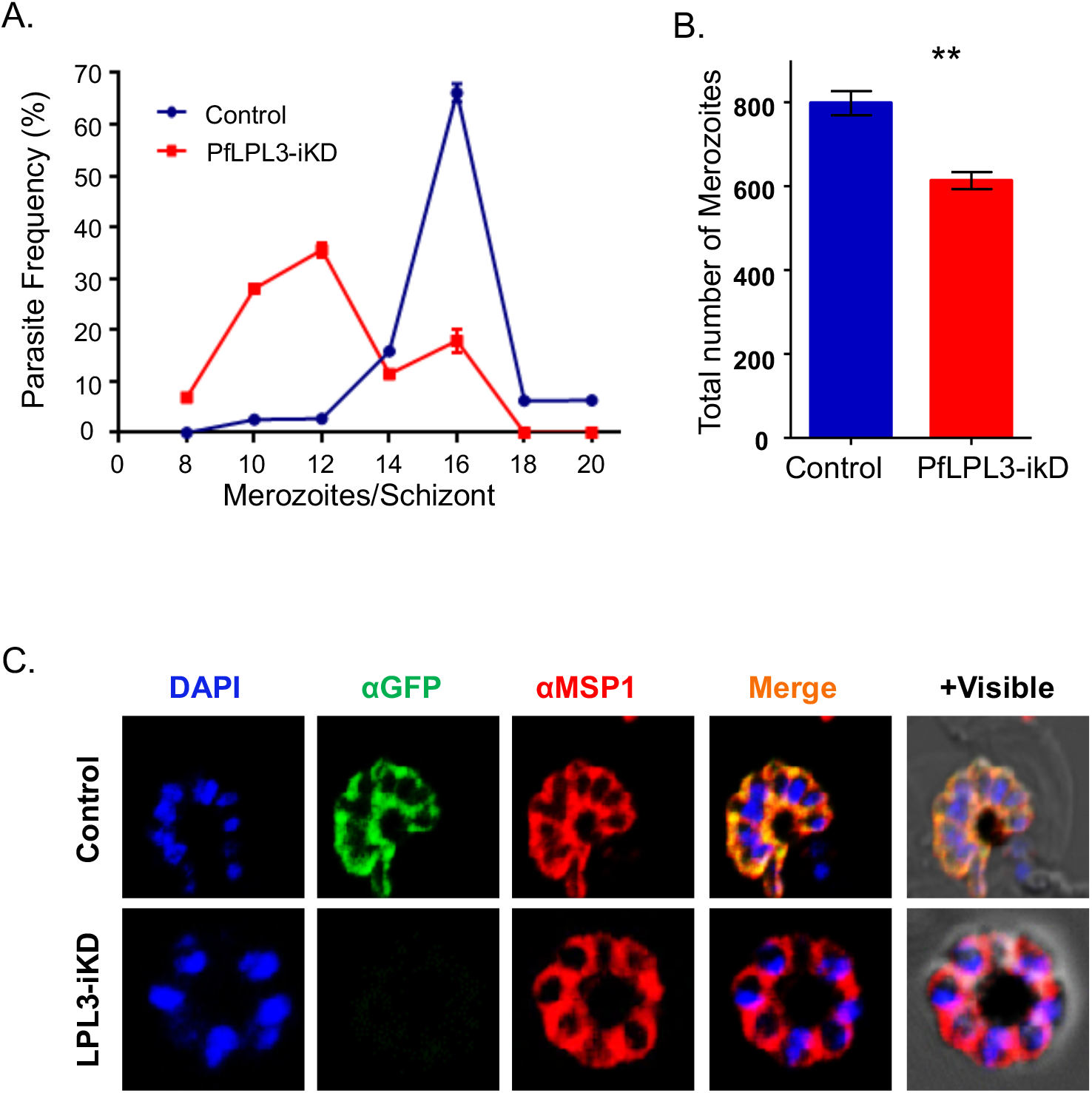
*Pf*LPL3 knock-down effect parasite schizogony: (A) Frequency distribution of number of merozoites per schizont in the control and *Pf*LPL3-iKD parasite cultures. (B) Graph showing reduction in the total number of merozoites in *Pf*LPL3-iKD parasite line in comparision to the control (*n*=50 segmented schizonts). ** *p* <0.01 (C) Fluroscent images of control and *Pf*LPL3-iKD parasites immunostained with anti-MSP1 antibody suggesting that there is no defect in the merozoites membrane in the *Pf*LPL3-iKD set.

### Downregulation of *Pf*LPL3 disrupts lipid homeostasis through decrease of neutral lipid content, concomitant increase of free fatty acids and phospholipids relative abundance

To assess the role of *Pf*LPL3 for parasite membrane biogenesis, we conducted exhaustive mass spectrometry-based lipidomic analyses on total lipid extracted from *Pf*LPL3-iKD set of parasites. Based on the predicted lysophospholipase function of *Pf*LPL3, i.e. releasing FA from existing (lyso)phospholipids, total FA content of parasites was initially assessed. Logically, the disruption of *Pf*LPL3 induced a significant reduction of total FA abundance (in nmol) (Fig. 5A) as quantified by lipidomic approach. Detailed analysis of the FA composition of total FA revealed significant reductions of the abundance of C16:0, C18:0 and C18:1 (Fig. 5B), all of which are known to be massively scavenged from the host (4)(12). We then sought to determine which lipid species were potentially affected the loss of *Pf*LPL3 and the reduction in FA content. We therefore analyzed the content of major FA-containing lipids, i.e. (a) neutral lipids making the bulk of lipid bodies/storage lipids: diacylglycerol (DAG) and triacylglycerol (TAG), (b) phospholipids (PL), major membrane lipids, and (c) free FA that are used as building blocks for both neutral lipids and phospholipids. The abundance of TAGs, DAGs, and PL was significantly lower in absence of *Pf*LPL3, whilst the abundance of FFA was significantly increased, together confirming the important role of *Pf*LPL3 for parasite membrane biogenesis (Fig. 5C). Importantly, the relative abundance profiles (in mol %) revealed that there was a significant decrease of TAGs and DAGs whereas FFA and phospholipids were significantly increasing in the absence of *Pf*LPL3 (Fig. 5 D1, 2, 3). This indicates that the downregulation of *Pf*LPL3 induces an accumulation of FFA, a decrease in the synthesis of TAGs and DAGs in favor of the synthesis of PL instead. Further TAG, DAG and PL composition analyses confirmed that it was the bulk of each lipid class that was impacted by the loss of *Pf*LPL3 rather than specific molecular species (Fig. S4), again emphasizing the central role of *Pf*LPL3 for parasite membrane homeostasis. Further analysis of the neutral lipid content shows that both the real and relative abundance of the TAG/DAG ratio is significantly reduced in the absence of *Pf*LPL3 (Fig. 5E), showing that a decrease in TAG following *Pf*LPL3 induces a concomitant decrease in parasite DAG content.

**Figure 5:**
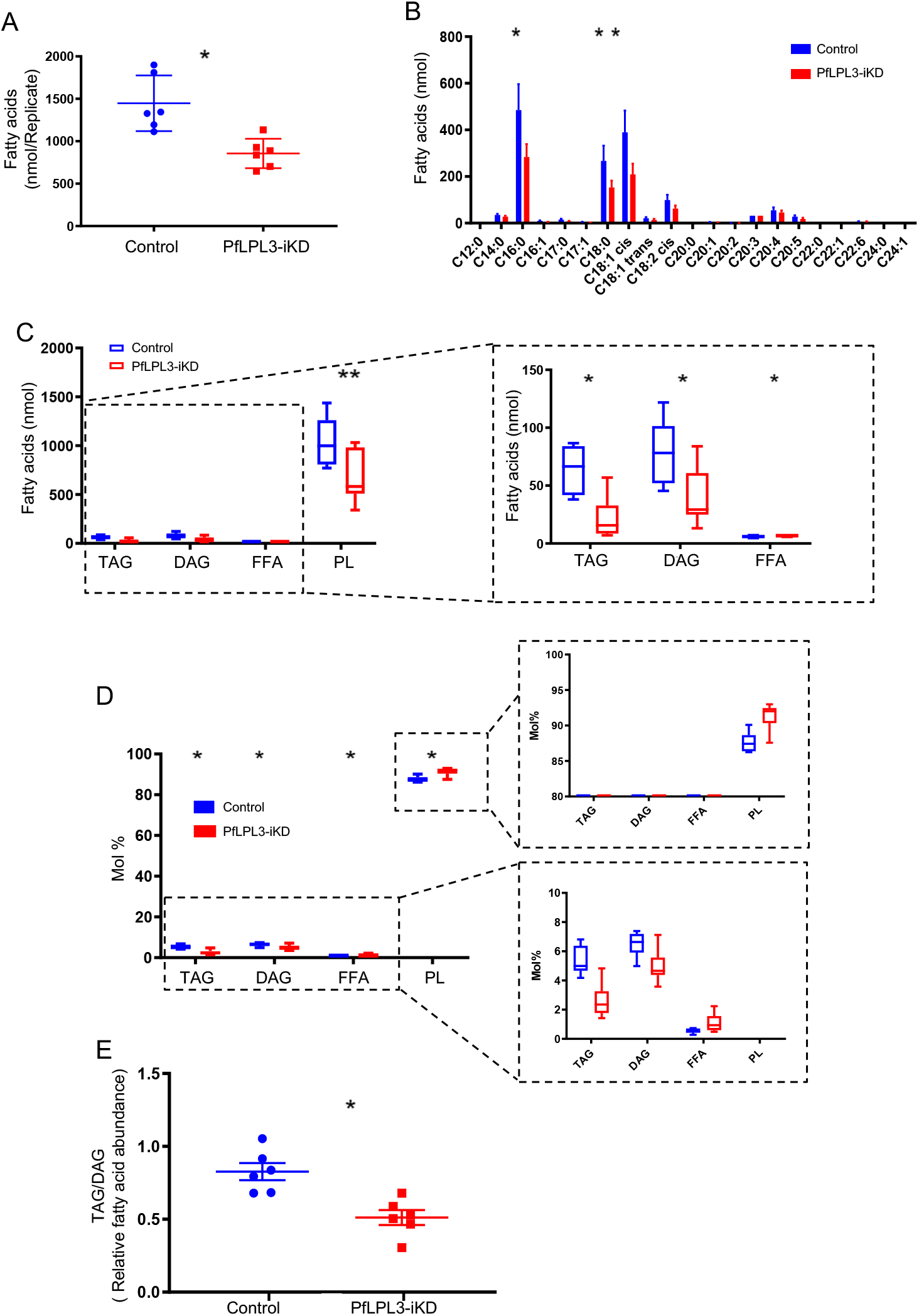
Lipidomic analyses reveal the role of *Pf*LPL3 to generate FAs and regulate DAG-TAG synthesis vs phospholipid synthesis: Synchronous transgenic parasites at ring stages were grown till late trophozoite stages (32h post infection) and lipid composition were assessed by mass spectrometry-based lipidomic analyses from *Pf*LPL3-iKD and control sets. Data is shown from six independent biological replicates. (A). Total fatty acid abundance (in nmol lipid/10^7parasites) reveal a significant decrease of total lipids in *Pf*LPL3-iKD. (B) Profile of fatty acid species in the 3D7 control and PfLPL3-iKD strains. There was a significant decrease in C160:0, C18:0 andC18:1 in the *Pf*LPL-iKD. (C) Quantification of the abundance (nmol) in TAG, DAG, FFA and PL between *Pf*LPL-iKD and control parasites revealed (D) Relative abundance of TAG, DAG, FFA, and PL showed significant decreases in TAG and DAG whereas there was a significant increase in FFA and PL. Right panels are enlarged sections of the left panel (E) Relative abundance of TAG to DAG ratio in *Pf*LPL3-iKD parasites as compared to control set. For all the assays P values are marked: * for *p* ≤ 0.05, and ** for *p* <0.005

### Developing a robust lysophospholipase enzymatic assay and identification of potent inhibitors against *Pf*LPL3

To determine the enzymatic activity of *Pf*LPL3, we expressed a recombinant protein corresponding to its lysophospholipase/hydrolase domain (41aa-338aa). The corresponding recombinant protein was purified, and which migrated on SDS-PAGE at the predicted size of ~70kDa (Fig. S5A). The purified recombinant *Pf*LPL3 was used to standardize a fluorescent-based activity assay to quantify its putative lysophospholipase activity, using LPC as a standard substrate (Fig. S5B). The recombinant *Pf*LPL3 showed concentration dependent lysophospholipase activity in the assay (Fig. S5C), confirming the predicted activity of *Pf*LPL3, and more specifically on LPC. The *K*m and *V*max values for the *Pf*LPL3 were found to be 49.25μM and 13113 nM/min/mg respectively (Fig. S5D). This result confirmed the lysophospholipase activity of the enzyme. The recombinant MBP (maltose binding protein) purified in same way as was used as a negative control and did not show any enzymatic activity. The *Z*-value for the activity assay was found to be ~0.9, showing the robustness of this activity assay, thus also suggesting it could be used for high throughput screening of compounds (Fig. S5E, F). Therefore, to identify putative inhibitors of *Pf*LPL3, we decided to screen the existing MMV antimalarial compounds, “Malaria Box”. Interestingly, two compounds MMV009015 and MMV665796, named as PCB6 and PCB7 respectively, were initially identified with a >70% inhibition of *Pf*LPL3 enzyme activity at 5μM concentration. We then determined the corresponding IC_50_ values of these compounds to be 1.26 μM and 0.92 μM respectively. Their *K*i were found to be 0.634 μM and 0.464 μM respectively (Fig. 6A)

**Figure 6:**
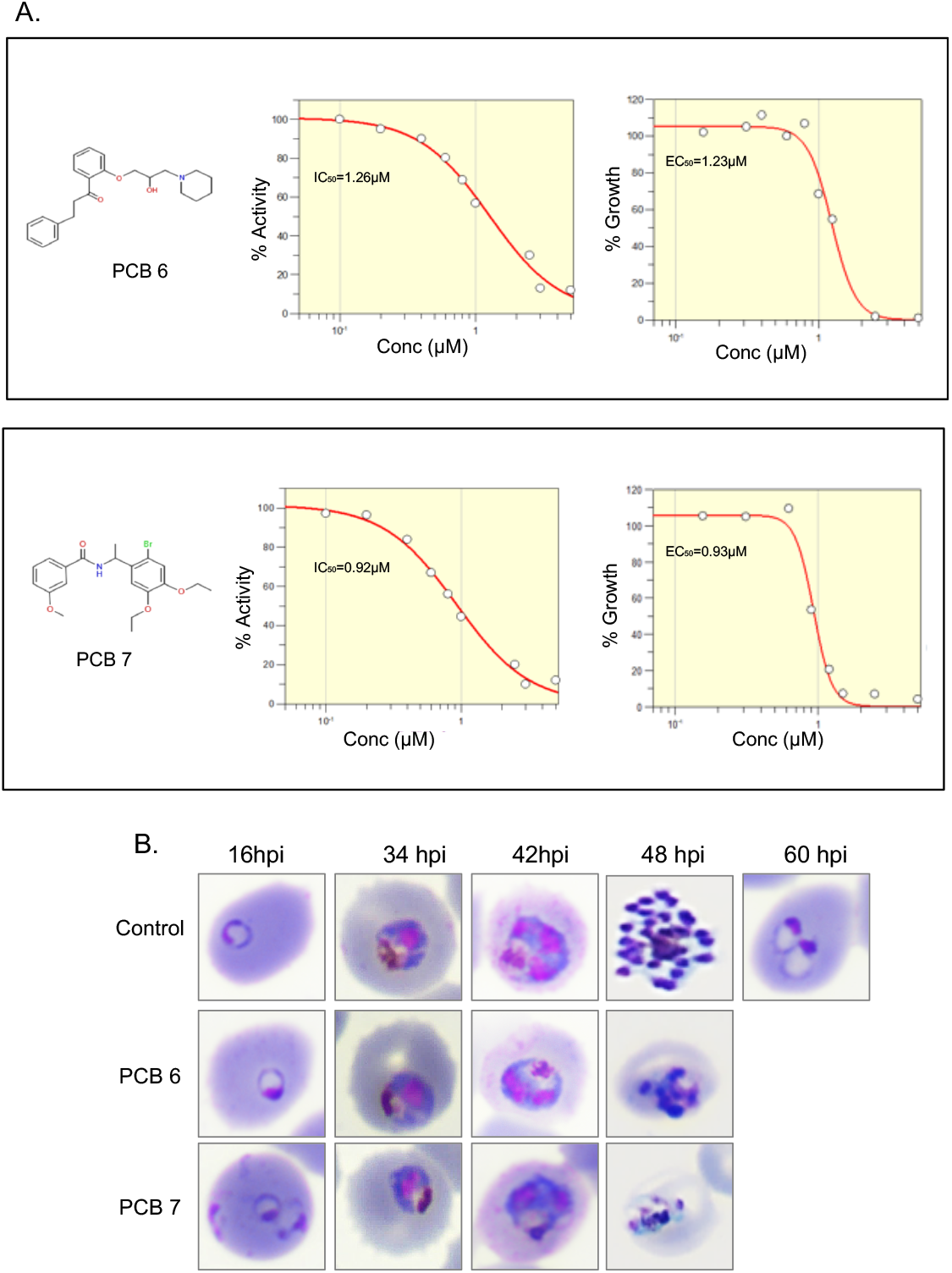
Identification of *Pf*LPL3 specific inhibitors from MMV Malaria Box: A robust *in vitro* activity assay was used to screen the ‘Malaria Box’ and potent inhibitors of *Pf*LPL3 are identified having parasiticidal efficacies. (A) Concentration dependent inhibition of enzymatic activity as well as inibition of asexual stage parasite growth by PCB6 (MMV009015) and, and PCB7 (MMV665796) compounds respectively. The IC_50_ values (enzyme inhibition) and EC_50_ values (parasite growth inhibition) for both compunds are indicated. (B) Effect on the morphology of *P. falciparum* parasites at different time points after treatment with *Pf*LPL3 inhibitors PCB6 and PCB7 at a concentration of EC_50_ values.

**Figure 7:**
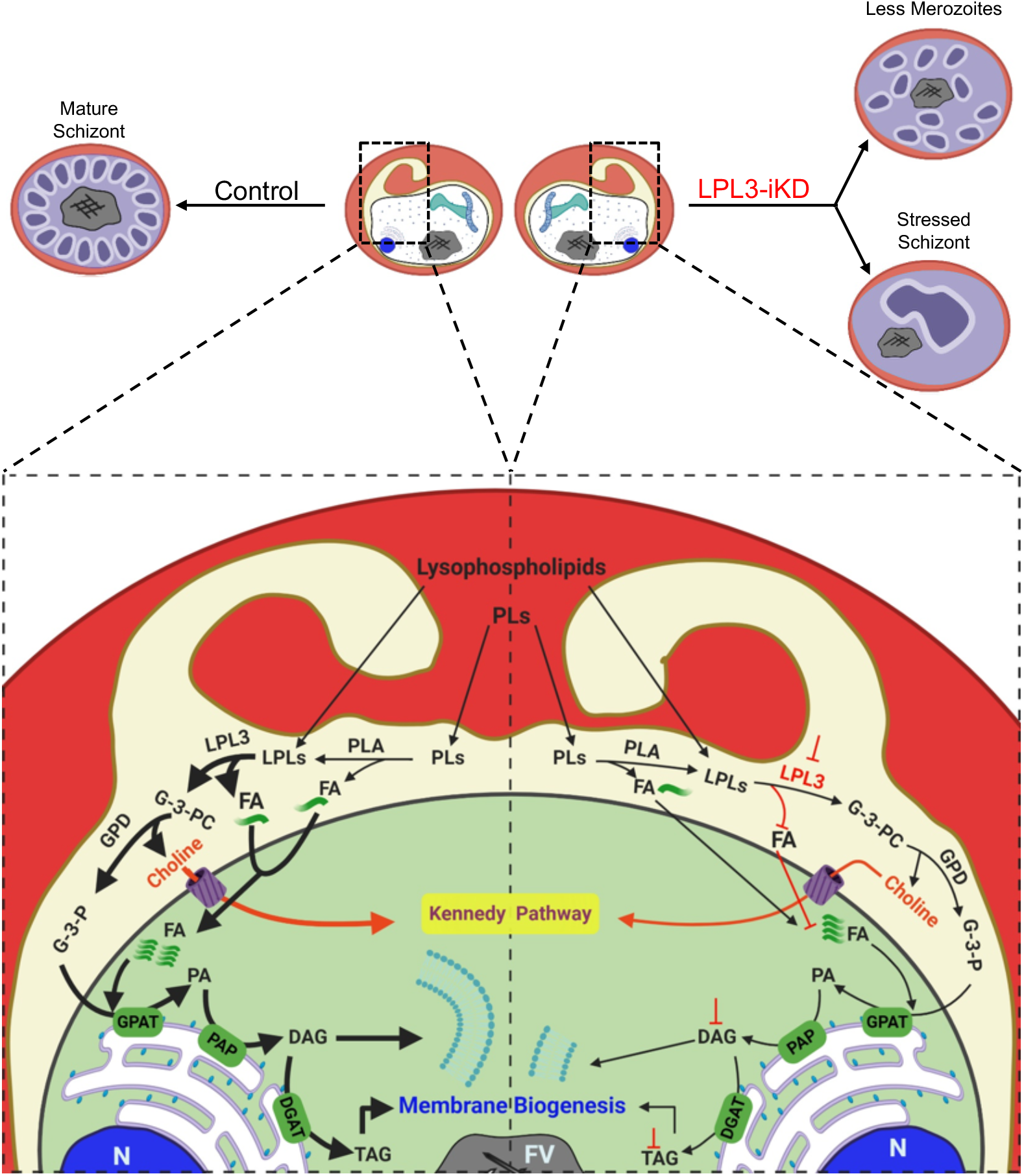
Model of *Pf*LPL3 function in *Plasmodium falciparum* asexual blood stage cycle: Schematic diagram showing metabolic pathways of lipid metabolism associated with proposed function of *Pf*LPL3. In the left panel, *Pf*LPL3 level is maintained in absence of glucosamine. At the late trophozoite stage, *Pf*LPL3 localizes in the PV of the parasite which cleaves the acyl chain from the lysophospholipids (acquired from the host or generated from phospholipid catabolism) results in generation of G-3-P and FAs. Which ultemately required for DAG and TAG synthesis. In presence of PfLPL3 neutral lipid and membrane biogenesis are normal which results normal schizogony. On the right panel, consequences of *Pf*LPL3 knockdown are shown. The decreased level of *Pf*LPL3 level in presence of glucosamine leads to desregulation of fatty acid content, which disrupts phospohlipid synthesis and development of new membranes. Therfore *Pf*LPL3 knock-down disruspt schizogony and cause reduction in number of merozoites in schizonts. PLs (Phospholipids), PLA (Phospholipase A), FA (Fatty acid), LPLs (Lysophospholipids), LPL3 (Lysophospholipase 3), G-3-PC (Glycero-3-Phosphocholine), G-3-P (Glycero-3-Phosphate), PA (Phosphatidic acid), DAG (Diacylglycerol), TAG (Triaclyglycerol), GPAT (Glycerol-3-phosphate acyltransferases), PAP (Phosphatidate phosphatase), DGAT (Diglyceride acyltransferase), FV (Food Vacuole), N (Nucleus).

### *Pf*LPL3 Inhibitors block *P. falciparum* schizont development

The two “Malaria Box” compounds PCB6 and PCB7 are known to show parasiticidal efficacies with EC_50_ at low μM range (Fig. 6A). We confirmed the EC_50_ of PCB6 and PCB7 compounds to be 1.26 μM and 0.9 μM respectively (Fig. 6A). Both inhibitors were assessed for their effect throughout parasite developmental cell cycle during asexual blood stages. It was found that treatment with compound PCB6 and PCB7 led to normal parasites development from ring to trophozoites/late-trophozoite, similar to the solvent control (Fig. 6B). However, more than 50% of these trophozoites were not able to fully develop into mature schizonts with their morphology appeared deeply affected. The morphological changes of parasites treated by PCB6 and PCB7 both timely coincided and phenocopied the morphological effect observed in *Pf*LPL3 knockdown (Fig. 6B). These results strongly indicate the specificity of these compounds for *Pf*LPL3. Both compounds therefore displayed promising anti-malarial activity and their mechanism of action likely targeting *Pf*LPL3 and its central function for parasite membrane biogenesis and progeny formation during schizont stages.

## Discussion

A number of studies highlight that metabolic pathways linked with lipid synthesis, catabolism and trafficking are essential for survival of malaria parasite (13)(14)(15, 16). During the intra-erythrocytic cycle of *P. falciparum* there is large increase in the phospholipid, neutral lipid and lipid-associated FA contents in the infected red blood cells (4, 6, 17). Indeed, the parasites require large amount of lipids to synthesise new membrane-bound organelles as well as for the assembly of new merozoites after cell division (18–20). To fulfill this massive requirement of lipids, the parasites *de novo* synthesize as well as scavenge lipids/phospholipids from host milieu during the erythrocytic cycle. Phospholipids play a central role in the progression of the life cycle of every parasite and its pathogenesis. Lysophospholipids, especially LPC, are highly present in the host cell external environment and are actively scavenged by intra-erythrocytic parasites. Importantly, LPC serve as major substrates for phospholipid synthesis via the Kennedy pathway through the generation of choline for further PC synthesis and concomitant de-acylation (i.e. loss of their FA chain) putatively catalyzed by action of unknown lysophospholipase (9) (8). The nature of the involved phospholipases and their exact function during parasite development are currently unknown. Interestingly, an unusual high number of LPLs have been identified in *P. falciparum* genome, however their putative roles, functions and localizations remain to be elucidated.

Here, to further understand lipid homeostasis and membrane biosynthesis in the parasite, we explored the role of a putative LPL in *P. falciparum, Pf*LPL3, during asexual life cycle of the parasite. *Pf*LPL3 is a member of lysophospholipase family in the parasite having α/β hydrolases domain and the GXSXG motif, which is the characteristic feature of lysophospholipases. We carried out detailed localization and functional studies to understand role of *Pf*LPL3 in parasite. A number of genetic tools have been developed for transient down regulation of target proteins in *Plasmodium* by tagging native genes (21–23). Here we have tagged the native gene at the C-terminus with GFP and a ribozyme system (10), which was used for both localization and transient knock-down of the target. *Pf*LPL3 was localized at the parasitophorous vacuole (PV) as well as in the tubulovesicular networks (TVN) extensions within host erythrocyte. The PV regions are at the interphase of host-parasite interaction during intracellular growth and are site of active transport of material from host to parasite through PVM. The PVM act as a molecular sieve for the parasite whereas TVN are extensions of PVM, which increases the surface area of the PVM to facilitate the more nutrient uptake and protein transport. Based on its putative function, the presence of *Pf*LPL3 in PV is quite intuitive and reveals that the PV/TVN regions are also sites for phospholipid uptake and metabolism. Glycerophosphodiesterase are enzymes that can act downstream of LPL and removes the choline moiety from glycerophospholipids. Recent study (24) shows tripartite distribution of glycerophosphodiesterase in the cytosol, PV and food vacuole. Presence of glycerophosphodiesterase as well as lysophospholipase, *Pf*LPL3, in the PV therefore of the parasite confirms this area as a major site for the catabolism of exogenously imported phospholipids into FA and respective head-group.

Transient down regulation of *Pf*LPL3 caused severe disruption in the development of trophozoites into schizonts stages. Trophozoite stage of the parasite is metabolically most active stage in which parasite generates and utilizes the available raw materials for organelle biogenesis, daughter merozoite cell formation, cytokinesis and karyokinesis; all these processes require extensive amount of lipids and membrane material. Those parasites in the *Pf*LPL3-iKD set which were able to develop from trophozoite into schizonts, were heavily compromised on number of merozoites developed, although these merozoites had properly developed membrane. In *P. berghei,* a phospholipase (PbPL) is also shown to be present in the PV and its depletion caused defect in the egress from the host hepatocytes (25). The *Pf*LPL3-iKD inhibited schizont development as well as reduced replicative fitness of parasites, which resulted in significant parasite growth inhibition.

Intracellular growth and segregation of *Plasmodium* requires lipid synthesis, which is essentially needed for new membrane development. Phospholipids are *de novo* synthesized/assembled by the parasite; however, their precursors (FA and polar heads) are scavenged from the host milieu. It has been shown that depletion in the serum LPC leads to decrease in the number of merozoites in rodent malaria model (8). Host scavenged LPC is essential for Phosphatidylcholine (PC) synthesis in the parasite; PC is generated by Kennedy pathway from cytidine diphosphate (CDP)-choline (26, 27), whereas in absence of choline, PC is generated via triple methylation of PE using ethanolamine and serine as external precursors. Our lipidomic analysis revealed that *Pf*LPL3 is pivotal for the generation of FA and subsequent lipid synthesis for parasite development. Specifically, we found that *Pf*LPL3 functions as a regulator to channel the high rate in FA influx from the host towards neutral lipid DAG and TAG synthesis. Its absence leads to an uncontrolled increase in phospholipid content, most likely blocking the normal development of trophozoite into schizonts and future merozoites. The neutral lipids play important role in eukaryotic cell cycle; DAG is known to activates protein kinase C by acting as second messenger (28), it also serves as a precursor for synthesis of TAG and major phospholipids. TAG is stored into cytosolic lipid droplets that detoxify and sequester FAs; whereas, TAG degradation by specific lipase can release these FAs to be utilized for membrane assembly (29). In asexual stage parasites, neutral lipids are stored in a lipid-body associated with the food-vacuole (30, 31). During parasite growth, hydrolysis of TAG can rapidly produce FAs to utilize in membrane synthesis (32). A general lipase inhibitor, Orlistat, was suggested to block utilization of stored TAGs, which inhibited parasite growth and caused developmental arrest in late stages (6). In similar way, reduction in TAG levels in *Pf*LPL3-iKD conditions, caused arrest of parasite in late stages, in addition, some of the parasite showed reduced number of merozoites, which points at hindrance in membrane biogenesis. Our data reveals that during blood stages, *P. falciparum* controls lipid synthesis and consequent formation of daughter cells by: (1) favoring massive generation and storage of FA building blocks through the synthesis of DAG and TAG; and (2) timely allowing the generation of structural phospholipids for merozoite formation through the utilization of FA from DAG and TAG, rather than constantly accumulating FFA and/or synthesize phospholipid from them, both of which being lipotoxic for the parasite.

One of the important motives to understanding the biology of the parasite is to identify functionally important metabolic pathways as drug targets, and subsequently design their specific inhibitors to develop as new antimalarials. Our study involving detailed localization, gene knock-down and lipidomic analyses clearly show that *Pf*LPL3 plays critical role in growth and segregation of the parasite during asexual cycle. We also showed that *Pf*LPL3 harbor enzymatic activity for lysophospholipase. Furthermore, a robust *in vitro* activity assay was established to screen the compound libraries. Medicine for Malaria Venture (MMV) established an open collaborative initiative in search of novel drugs for neglected tropical diseases. Based upon extensive screening strategy, they identified a set of drug-like compounds having anti-malarial activity, and labelled this set as ‘Malaria Box’(33). These compounds are new leads to develop further. However, there is need to identify potential targets for these compounds in the parasite, which may help medicinal chemistry approaches to design diversity library and identify potent antimalarial. Screening of the “Malaria Box” by using the *Pf*LPL3 activity assay identified two hit compounds, labelled as PCB6 and PCB7; both the compounds have parasiticidal efficacy and block the schizont development in *P. falciparum,* which phenocopies the gene knock-down phenotype, and thus consistently confirms the role of *Pf*LPL3. Further, both the compounds showed very close efficacies to inhibit enzyme activity and parasite growth, IC_50_ and EC_50_ respectively, in the low micromolar range. These results suggest that both the compounds specifically targeting the *Pf*LPL3 in the parasite. Hence, these can be further developed through medicinal chemistry campaign to develop potent antimalarials, in addition of being specific molecular tools to study lipid synthesis and membrane biogenesis during schizogony and merozoite formation.

## Materials and Methods

### Parasite culture, plasmid construct and parasite transfection

*Plasmodium falciparum* strain 3D7 was cultured in RPMI media (Invitrogen) supplemented with 0.5% (w/v) Albumax™ (Invitrogen) 4% haematocrit. Culture was kept static in a gas mixture of 5% carbon dioxide, 5% oxygen and 90% nitrogen at 37°C. To generate *Pf*LPL3_GFP_*glm*S construct, a C-terminal fragment (702 base pair) of *pfLPL3* gene was amplified using specific primers 1222A (gcACTAGTAGTATTTTAGAATCTGAAACG) and 1223A (gcGGTACCTTTCTTTTCTTTTTCTTTTTGC) and cloned in to GFP_*glm*S vector. The C-terminal fragment was cloned in frame to the N-terminus of GFP in the *Spe*I and *Kpn*I restriction enzymes sites. Parasite cultures were synchronized by repeated sorbitol treatment and 100 μg of purified plasmid DNA (Plasmid Midi Kit, Qiagen, Valencia, CA) was transfected in *P. falciparum* by electroporation (310 V, 950 μF) (Crabb et. al. 2004). Transfected parasites were selected over 2.5 μg/ml blasticidin (Calbiochem) and subsequently subjected to on and off cycling of blasticidin drug to promote integration of the plasmid in the main genome. Integration was confirmed by PCR using 1270A and 1234A and by western blotting also by using anti-GFP antibody.

### Conditional knock-down and in-vitro growth assays analysis

To assess the effect of knock-down of *Pf*LPL3 on parasite, *Pf*LPL3_ GFP_*glm*S transgenic parasites were tightly synchronized with 5% sorbitol and different concentration (0mM, 1.25mM, 2.5mM, and 10mM) of glucosamine (Sigma-Aldrich) was added in 24 well plates containing 4% hematocrit and 1% parasitemia. For microscopic analysis and morphology of the parasites, Giemsa stained smears were prepared at every 8 hr interval, from both glucosamine treated and control sets. Parasite growth was assessed at 48 and 96 hours after the addition of glucosamine by flow cytometry using BD FACS Calibur (Beckton Dickinson); briefly, cells were incubated with EtBr for 30 minutes at 37 C in dark and after two washing with PBS, 100000 events were acquired using BD FACS Calibur system (Beckton Dickinson). To assess the effect of glucosamine on the down regulation of *Pf*LPL3 at protein level, *Pf*LPL3_GFP_*glm*S ring stage parasites were grown in presence of various concentration of glucosamine (1.25mM, 2.5mM, 5mM, 10mM), harvested at schizonts stage and subjected to the western blotting.

### Parasite fractionation and Western blot analysis

Parasites were harvested at trophozoite stage using 0.15% saponin for RBC lysis, the supernatant was collected, and parasites pellet was lysed by freeze-thaw cycle. Laemmli buffer was added to both the fractions and proteins were separated in 12% SDS-PAGE; the proteins were transferred to PVDF membrane (Millipore) and incubated with blocking buffer (4 % skim milk in 1 × PBS) followed by incubation with the primary antibody: (monoclonal anti-GFP mice 1:5000 (Roche), rabbit anti-GFP 1:10000, rabbit anti-BiP 1:15000 or monoclonal anti-spectrin 1:1000). Blots were washed 5 times with 1 PBS, probed with HRP conjugated secondary antibody (1:100000) and visualized using ECL detection kit (Thermo-scientific).

### Immuno-fluorescence assay (IFA) and fluorescent microscopy

The parasites were fixed in 4% paraformaldehyde. After permeabilization with 0.1 % TritonX-100, 10% bovine serum was used for blocking; cells were incubated with rabbit polyclonal anti-GFP (1:500) or mice polyclonal anti-SERA5 (1:500) for 2 hours and subsequently with Alexa Flour-488 and- 594 labelled secondary anti-body. Labeled parasites were washed three times with 1 × PBS. Parasite nuclei were stained with DAPI at a final concentration of 5 μg/ml. The membrane structures in parasitized erythrocytes were labelled with BODIPY TR-ceramide (Invitrogen) with a final concentration of 1μM. Images were captured by Nikon A1 confocal laser scanning microscope and analysed by Nikon-nis element software (version 4.1). The 3D images were constructed by using series of Z-stack images and IMARIS 7.0 (Bitplane Scientific) software.

### Lipid extraction from *P. falciparum* and liquid chromatography-mass spectrometry analysis

Total lipids were extracted from treated (*Pf*LPL3-iKD) and control parasites. First parasites (4 × 10^8^ cell equivalents) were harvested by using 0.15% saponin treatment. The total lipid samples were spiked with 20 nmol C21:0 phosphatidylcholine, and extracted by chloroform: methanol, 1:2(v/v) and chloroform: methanol, 2:1 (v/v). The pooled organic phase was subjected to biphasic separation by adding 0.1% KCl and was then dried under N2 gas flux prior to being dissolved in 1-butanol. For the total fatty acid analysis, an aliquot of the lipid extract was derivatized on-line using MethPrep II (Alltech) and the resulting FA methyl esters were analyzed by GC-MS as previously described. For the quantification of each lipid, total lipid was separated by 2D HPTLC using chloroform/methanol/28% NH4OH, 60:35:8 (v/v/v) as the 1st dimension solvent system and chloroform/acetone/methanol/acetic acid/water, 50:20:10:13:5 (v/v/v/v/v) as the 2nd dimension solvent system. Each lipid spot was extracted for quantification of fatty acids by gas chromatography-mass spectrometry (Agilent 5977A-7890B) after methanolysis. Fatty acid methyl esters were identified by their mass spectrum and retention time and quantified by Mass Hunter Quantification Software (Agilent) and the calibration curve generated with fatty acid methyl esters standards mix (Sigma CRM47885). Then each lipid content was normalized according to the parasite cell number and a C21:0 internal standard (Avanti Polar lipids). All analyses were performed in triplicate or more (*n*=3). The *p* values of ≤ 0.05 from statistical analyses (Student’s *t*-test) obtained from GraphPad software analysis were considered statistically significant.

### Cloning, Expression and purification of recombinant protein

Hydrolase domain of *Pf*LPL3 was amplified by using specific primers 1270A (gcGGATCCACCATGGAAGTATCATAATATTAATACATGGGTTAGC) and 1271A (gcCCTAGGTCCTTTTTCGATTGTTA TAACATGAC) and cloned in to pETM41 vector between *Nco*I and *Xho*I sites to give pETM41-His6-MBP-PfLPL3 construct. The recombinant protein *Pf*LPL3 was expressed as soluble protein in the cytosol of the *E. coli* BL21(DE3) codon^+^ cells. For expression, bacterial cells having pETM41-His6-MBP-PfLPL3 was grown in of LB medium supplemented with kanamycin at 37°C to an absorbance (A600) of ~0.8-1.0. Expression of the fusion proteins was induced using 1mM isopropyl β-D-thiogalactoside (IPTG) overnight at 16°C. Next day cells were harvested and resuspended into resuspension buffer (25mM Tris, pH7.4, 500mM NaCl, 10mM Imidazole) and lysed by sonication (Vibra cell sonicator). The supernatant was subjected a combination of Ni-NTA affinity chromatography followed by amylose affinity chromatography. Eluted fractions were subjected to SDS PAGE and western blot analysis to assess the purity of the purified recombinant protein.

### Lysophospholipase (LPL) activity assay and enzyme kinetics

For assessing the enzymatic activity of the recombinant *Pf*LPL3, a lysophospholipase (LPL) activity assay was established using lysophosphatidylcholine as substrate. The activity assay reaction mixture (200 μl total volume) contained 16:0 lysophosphatidylcholine (LPC) as substrate, 20μg (285pmol) of recombinant protein, 0.1 unit glycerophosphodiesterase (Sigma), 0.2 U/ml choline oxidase (Sigma 26978), 2U/ml horseradish peroxidase (Sigma P8125), 100μM Amplex Red (Invitrogen) in a reaction buffer (50mM Tris pH 8.0, 5mM CaCl_2_). The *Pf*LPL3 cleaves the acyl chain from LPC resulting the generation of glycerophosphocholine on which glycerophosphodiesterase acts and cleaves the choline moiety. This choline was oxidized by choline oxidase to produce betaine and H_2_O_2_. Finally, H_2_O_2_ in presence of horseradish peroxidase, reacts with Amplex Red reagent in 1:1 stoichiometry to generate the highly fluorescent product resorufin, which has absorption and fluorescence emission maxima of approximately 571 nM and 585 nM respectively. The LPL activity was monitored by measuring fluorescence intensities (530 ex/590 em) for 6 hrs using spectramax M2 microplate reader. The assay was performed by varying the amount of protein and substrate to get optimum LPL activity. The kinetic constant *K*m and *V*max were determined by fitting Michalies-Menten curve using the Graph Pad Prism V5.0 software package.

### Standardization of a robust lysophospholipase (LPL) activity assay and screening of compound library

Robustness of the activity assays was assessed by conducting HML test for a period of two days in accordance for statistical test to calculate the Z-value. The HML assay employed three different reaction wells: **<u>H</u>**igh activity well i.e. lysophospholipase activity well with 20μg enzyme; **<u>M</u>**edium activity well i.e. in the presence reduced amount of enzyme, (10μg protein); and the **<u>L</u>**ow activity well that is the substrate control well without the enzyme; this set of three wells (HML) was repeated throughout the 96-well plate and the assay was repeated for two consecutive days. The Z-value was calculated based on the variations in activity between the wells/plates (34).

To identify the *Pf*LPL3 targeting antimalarial compounds, “Malaria Box” compound library obtained from Medicines for Malaria Venture (MMV, Switzerland), was screened using the standardized *Pf*LPL3 activity assay; briefly, the recombinant enzyme (20μg) was incubated with different concentrations of each of the compound or DMSO alone; the reactions were initiated by addition of the activity assay reaction mixture to a final volume of 200μl and the substrate hydrolysis was monitored as described above. To assess the effect of selected *Pf*LPL3 inhibitors on parasite growth and morphology, synchronized parasite cultures at 1% ring stage parasites were treated with varied concentrations of each of the selected compounds (5.0 to 0.2μM), parasite growth was estimated in next cycle by flow-cytometry. DMSO was used as a control.

### Statistical Analysis

The data sets were analysed using GraphPad Prism ver 5.0 to calculate *K*m, *V*max, IC_50_ and EC_50_ values, and the data were compared using unpaired Student’s *t*-test.

## Acknowledgements

We are grateful to Philip J. Shaw for providing vector pGFP_*glm*S and useful suggestions. We thank Rotary blood bank, New Delhi for providing the RBCs. PKS is supported by research fellowship from University Grant Commission, Govt. of India. VT is supported by BioCARe Women Scientist fellowship from Department of Biotechnology, Govt. of India. The research work in AM’s laboratory is supported by Centre Of Excellence grant (BT/COE/34/SP15138/2015) Flagship Project grant (BT/IC-06/003/91) from Department of Biotechnology, Govt. of India. CYB and YYB are supported by Agence Nationale de la Recherche, France (Grant ANR-12-PDOC-0028-Project Apicolipid), Finovi programs, LABEX ParaFrap, France (grant number ANR-11-LABX-0024), the Université Grenoble Alpes (IDEX ISP) Apicolipid) and Région Auvergne Rhone-Alpes for the lipidomics analyses platform (Grant IRICE Project GEMELI). The research work was supported by Indo-French Collaborative Research Program Grant (Project 6003-1) to AM and CYB by the CEFIPRA. The funders had no role in study design, data collection and interpretation, or the decision to submit the work for publication.

## Supplementry Figures

**Figure S1:**
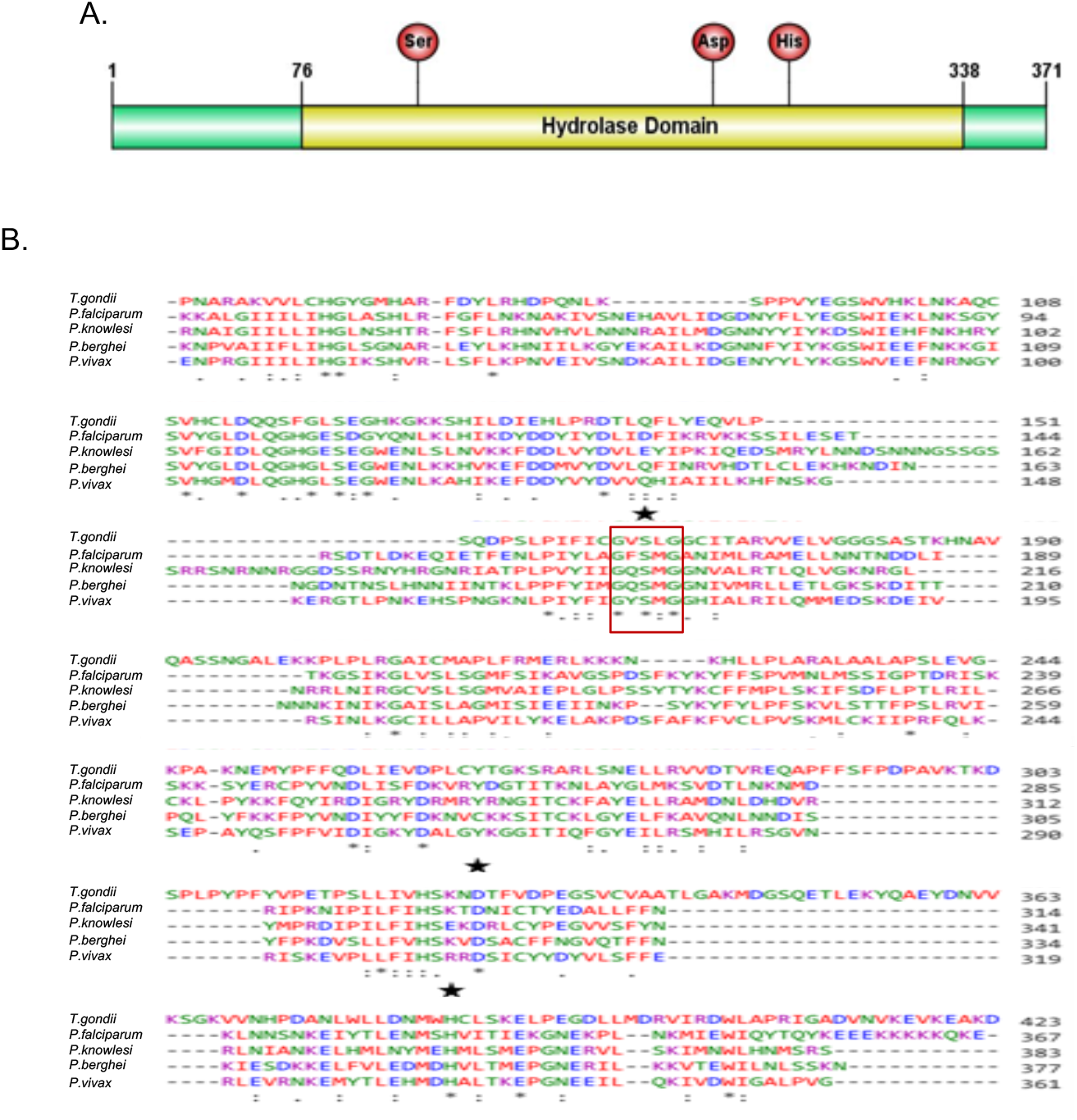
Domain organisation and multiple sequence alignment of *Plasmodium* LPL3 homologues: (A) Schematic diagram of *Pf*LPL3 showing 262 amino acids long hydrolase domain, location of active site residues is marked. (B) Multiple amino acid sequence alignment of *Pf*LPL3 with its homologs in other organisms (*T. gondii, P. knowlesi, P. berghei, P. vivax*) showing the characteristic motif and active site residues are conserved among these proteins. Red boxes indicate the conserved motifs and asterisk indicates the catalytic residues.

**Figure S2:**
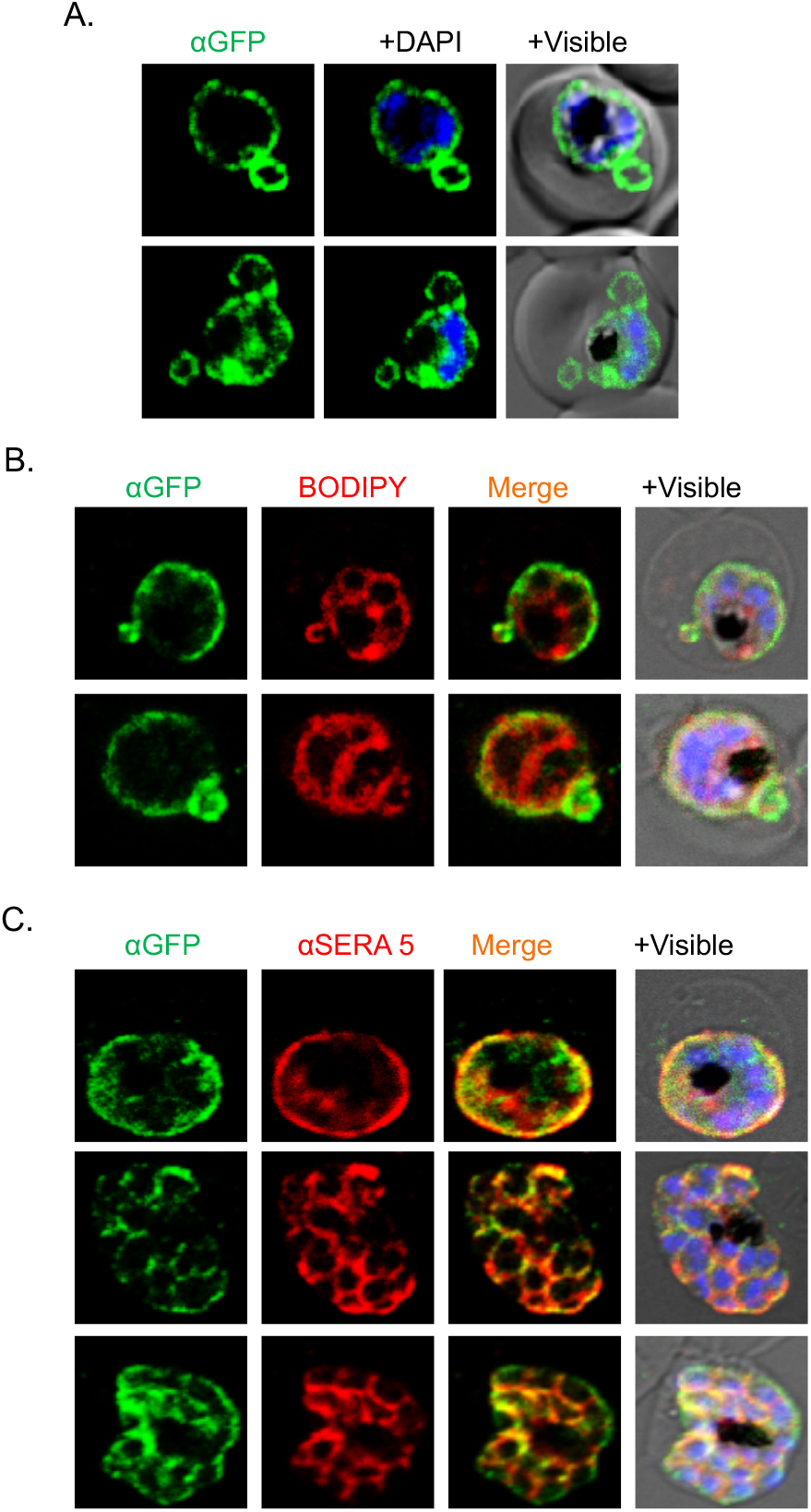
Co-localization of *Pf*LPL3 with PV and PVM markers: (A) Fluorescent microscopy image of the transgenic parasites shows that the GFP-fusion protein present in the PV and tubulovesicular network (TVN) region. (B) Fluorescent microscopic images of transgenic parasites stained with mambrane probe BODIPY-TR ceramide, showing GFP fusion protein at the parasite periphery in late parsite stages. (C) Fluorescent microscopic images of trangenic parasites immunostained with anti-SERA5 antibody showing colocalisation of *Pf*LPL3 with SERA5 in PV region. Parasite nuclei were stained with DAPI and images were acquired by confocal laser scanning microscope.

**Figure S3:**
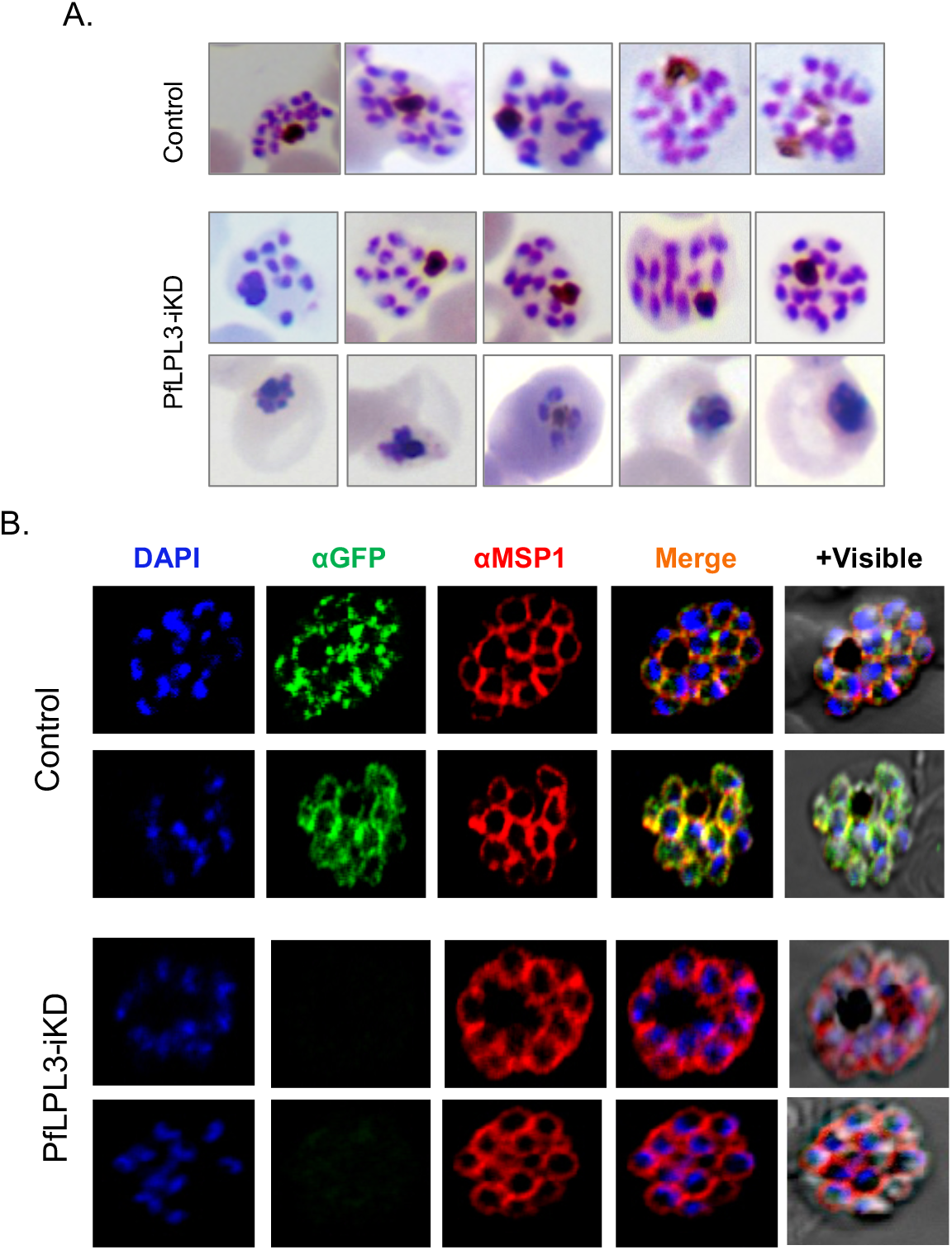
(A) Schizont development and merozoite seggregation is hindered in *Pf*LPL3-iKD parasites. Giemsa stained smears from the *Pf*LPL3 knock-down parasites set showing that the glucosamine treated parasites have stressed parasites as well as reduced number of merozoites per schizont as compared to control set which has mature schizonts with higher number of merozoites per schizonts. (B) Effect of *Pf*LPL3 knock down on merozoite membrane. Fluorescent images of the wild type and the *Pf*LPL3-iKD parasites immuno-stained with anti-MSP1 antibody. Although the number of merozoites were reduced in *Pf*LPL3-iKD, these merozoites have intact plasma membrane as in case of control. Parasite nuclei were stained with DAPI and images were acquired by confocal laser scanning microscope.

**Figure S4:**
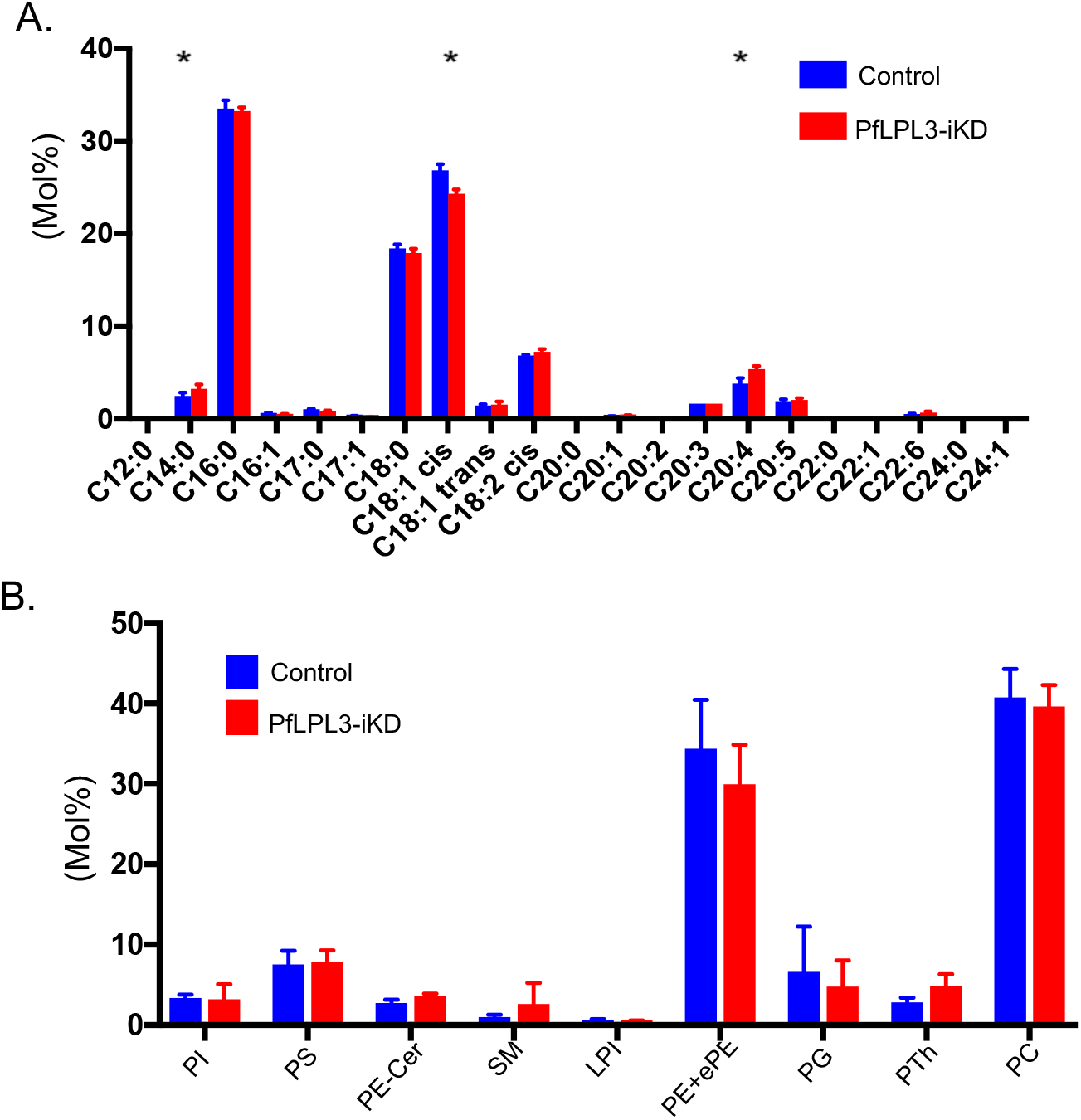
Alteration in lipid composition by inducible knock-down of *Pf*LPL3 in the parasites: (A) Graphs showing compsition (Mol%) of different clsasses of fatty acids, there was no alteration in abundance in different fatty acids classes between *Pf*LPL3-iKD and control. (B) Graphs showing compsition (Mol%) of different clsasses of phosopholipids, there was no alteration in abundance of different classes except minor changes in PE.

**Figure S5:**
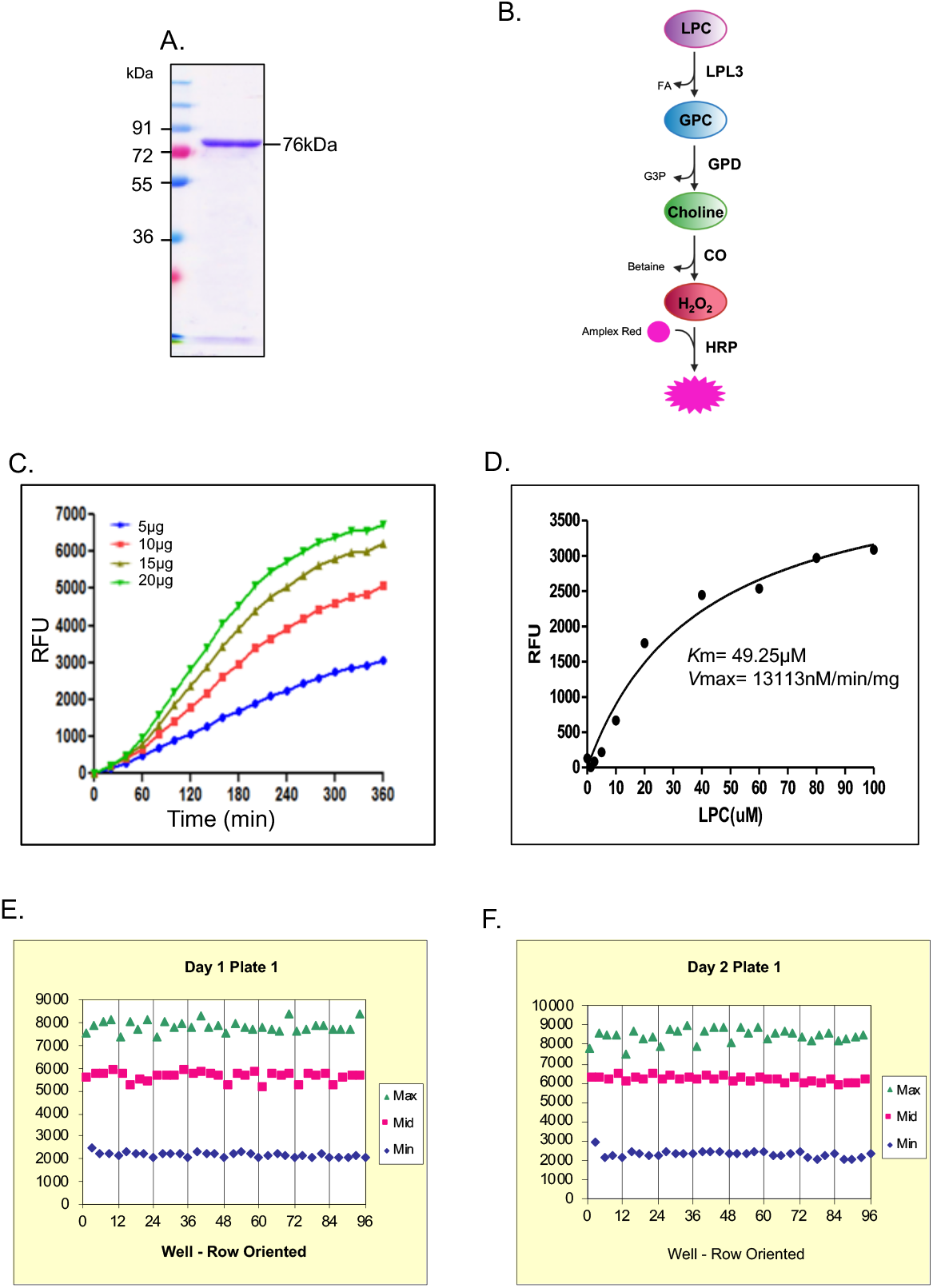
Standardization of lysophospholipase specific activity of the recombinant *Pf*LPL3 for screening of MMV ‘Malaria Box’ compound library: (A) Coomassie stained SDS-PAGE showing purified recombinant *Pf*LPL3. (B) Representative diagram of reaction and components involved in the LPL activity assay. LPC was used as a substrate The *Pf*LPL3 cleaves the acyl chain from LPC resulting the generation of glycerophosphocholine on which glycerophosphodiesterase acts and cleaves the choline moiety. This choline was oxidized by choline oxidase to produce betaine and H_2_O_2_. Finally, H_2_O_2_ in presence of horseradish peroxidase, reacts with Amplex Red reagent in 1:1 stoichiometry to generate the highly fluorescent product resorufin, and fluorescence of resorufin was measured. (C) Graphical representation of LPL activity of *Pf*LPL3 in presence of varying amount of protein. (D) Line graph of Michelis-Menten fit for *Pf*LPL3 activity in presence of the 20μg (85pmoles) of recombinant protein. The *K*m and *V*max values of *Pf*LPL3 were found to be 49.25μM and 13113nM/min/mg respectively. (E) & (F) Robustness of the activity assay was assessed by estimating Z-value carried out in HML format in 96-well plate for two consecutive days, the Z-value was found to be ~0.9, suggesting robustness of the assay. The screening was carried out at 5μM concentration, two compounds were identified to inhibit >70% enzyme activity which are labelled as PCB6 and PCB7.

## Notes

### Competing Interest Statement

The authors have declared no competing interest.

